# Finding differentially expressed sRNA-Seq regions with srnadiff

**DOI:** 10.1101/768838

**Authors:** Matthias Zytnicki, Ignacio González

**Affiliations:** INRA, MIAT, Toulouse, Castanet 31 326 France

## Abstract

Small RNAs (sRNAs) encompass a great variety of different molecules of different kinds, such as micro RNAs, small interfering RNAs, Piwi-associated RNA, among other. These sRNA have a wide range of activities, which include gene regulation, protection against virus, transposable element silencing, and have been identified as a key actor to study and understand the development of the cell. Small RNA sequencing is thus routinely used to assess the expression of the diversity of sRNAs, usually in the context of differentially expression, where two conditions are compared. Many tools have been presented to detect differentially expressed micro RNAs, because they are well documented, and the associated genes are well defined. However, tools are lacking to detect other types of sRNAs, which are less studied, and have an imprecise “gene” structure. We present here a new method, called srnadiff, to find all kinds of differentially expressed sRNAs. To the extent of our knowledge, srnadiff is the first tool that detects differentially expressed sRNAs without the use of external information, such as genomic annotation or reference sequence of sRNAs.

**Author summary:** We present here a new method for the *ab initio* discovery of differentially expressed small RNAs. The standard method, sometimes named *annotate-then-identify*, first finds possible genes, and tests for differential expression. In contrast, our method skips the first step and scans the genome for potential differentially expressed regions (the *identify-then-annotate* strategy). Since our method is the first one to use the *identify-then-annotate* strategy on sRNAs, we compared our method against a similar method, developed for long RNAs (derfinder), and to the *annotate-then-identify* strategy, where the sRNAs have been identified beforehand using a segmentation tool, on three published datasets, and a simulated one. Results show that srnadiff gives much better results than derfinder, and is also better than the *annotate-then-identify* strategy on many aspects. srnadiff is available as a Bioconductor package, together with a detailed manual: https://bioconductor.org/packages/release/bioc/html/srnadiff.html

## Introduction

The eukaryotic small RNA (sRNA) repertoire has been greatly enriched by the use of small RNA sequencing, which provides a wide range of small RNAs (usually defined as RNAs with size not greater than 200) of a set of given cells. These sRNAs include the well-known micro RNAs, but also tRNA-derived RNA fragments (tRFs), small interfering RNAs (siRNAs), Piwi-associated RNAs (piRNAs), among others [1, 2]. These sRNAs are involved in many stages of development and diseases, in genetic and epigenetic pathways. The interest in these small molecules has been constantly growing.

The role of these sRNAs is usually understood *via* a differential expression protocol, *e.g.* healthy *vs* sick, or wild type *vs* mutant. Most analyses only focus on miRNAs, partly because miRNAs genes (pre-miRNAs or mature miRNAs) are known, or can be efficiently discovered. These genes are then used in the standard (messenger) RNA-Seq protocol for differential expression: reads are mapped to the reference genome, gene expression is quantified using read counts, and differential expression is tested.

While this approach can be used for other sRNAs, such as tRFs (because they originated from tRNAs), it is not directly applicable to all sRNAs, such as siRNAs or piRNAs, because the corresponding “genes” are not known (and possibly even not defined). Finding boundaries of sRNAs based on the expression profile is also a difficult task, because the expression profile of the sRNAs can be very diverse: miRNAs and tRNAs, for instance, exhibit sharp peaks, whose sizes are approximately the size of a read. The expression profiles of siRNAs and piRNAs are usually wider, ranging possibly several kilo-bases or more, with an extremely irregular contour. These sRNAs can even be found in clusters, and the aggregation of several, possibly very different, profiles, makes it hard to discriminate them.

So far, current researchers have three options. First, the reads can be mapped to a set of reference sequences, such as miRBase [3], which stores all the known miRNAs. The counts are then stored in a matrix, where the rows are the features (here, the miRNAs), the columns are the samples, and the cells are the number of reads that match a given feature, in a given sample. This method has been used by [4], who analyzed the sRNA-Seq data of lung tumors compared to adjacent normal tissues. This method can obviously only find features that are previous detected, and usually restrict the analysis to only one, or few, classes of small RNAs.

The second option has been used by an other article, that reused the previous dataset in order to find differentially expressed snoRNAs and piRNAs [5]. Here, the authors mapped the reads to the genome, and compared the mapped reads with external annotations, here piRNAs (from piRNABank [6]), and snoRNAs (from UCSC genome browser annotation [7]). The authors claimed that the approach is more exhaustive, since —especially in human— the annotation files which are provided by the existing repositories include a wide diversity of small RNAs. However, the analysis still restrict to known and annotatated small RNAs.

Some popular tools for sRNA differential expression, such as UEA sRNA Workbench [8] and sRNAtoolBox [9] use a combination of these methods. While these method works fine for miRNAs, and other well-know sRNAs, they cannot detect other types of sRNAs, or be applied to an unannotated genome.

Other methods, such as BlockClust [10], SeqCluster [11], or ShortStack [12], include a clustering step, which assemble the reads into longer transcripts. Then, the user can proceed to the standard messenger RNA-Seq pipe-line: counting reads that co-localize with each transcript, and testing for differential expression. These tools may or may not use an annotation file. The downside of this approach is that it requires significantly more work and time to cluster the reads into transcripts.

Recently, [13] presented derfinder, a new method for discovering differentially expressed (long) genes. Briefly, the authors find differentially expressed regions at the nucleotide resolution, regardless of the annotation. This promising method, however, has been designed for RNA-Seq and works poorly on sRNA-Seq, because sRNA expression profiles are very different from the longer, somewhat uniform, expression profiles of the exons.

In this work, we present a new method, srnadiff, that finds differently expressed small RNAs, using RNA-Seq data alone, without annotation. We show that srnadiff is more efficient than other methods, and that it detects a wide range of differentially expressed small RNAs.

## Materials and methods

### Description of the method

The method is divided into two main steps, which are described hereafter. The outline of the method is given in Fig 1.

**Fig 1.**
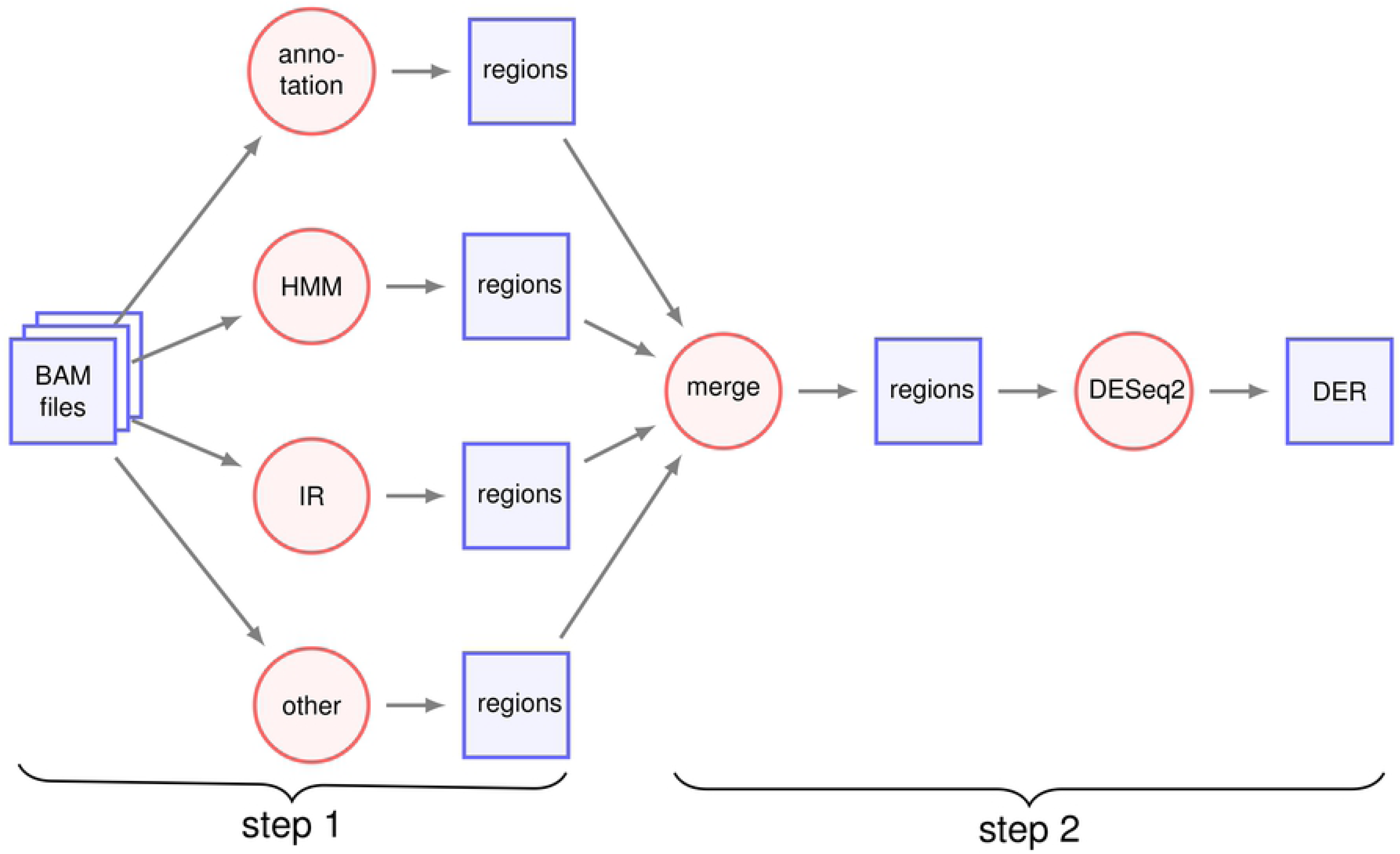
Outline of the method. Files and intermediate data are displayed in blue rectangles, and algorithms are displayed in red circles. DER: differentially expressed regions.

#### Step 1: find candidate regions

In this step, several methods are used to produce genomic intervals that are potential differentially expressed regions. We implemented three methods: naïve, HMM, and IR (described in the next sections), but, in principle, any method can be added.

#### Step 2: merge regions

The intervals provided by the previous step may overlap since several methods may give similar intervals. The aim here is to keep only the best non-overlapping regions.

To do so, the intervals provided in the previous steps are used as standard genes and we use the RNA-Seq standard pipe-line.

- The expression of the intervals is quantified for each condition (a read is counted for every interval it overlaps).
- DESeq2 is used to compute a p-value for each intervals.

When two regions, *i*_1_ and *i*_2_, overlap, *i*_1_ *dominates i*_2_ iff its p-value is less than the p-value of *i*_2_. A first possibility is to give undominated intervals to the user, but we found that it removes many interesting intervals (see Fig 2).

**Fig 2.**
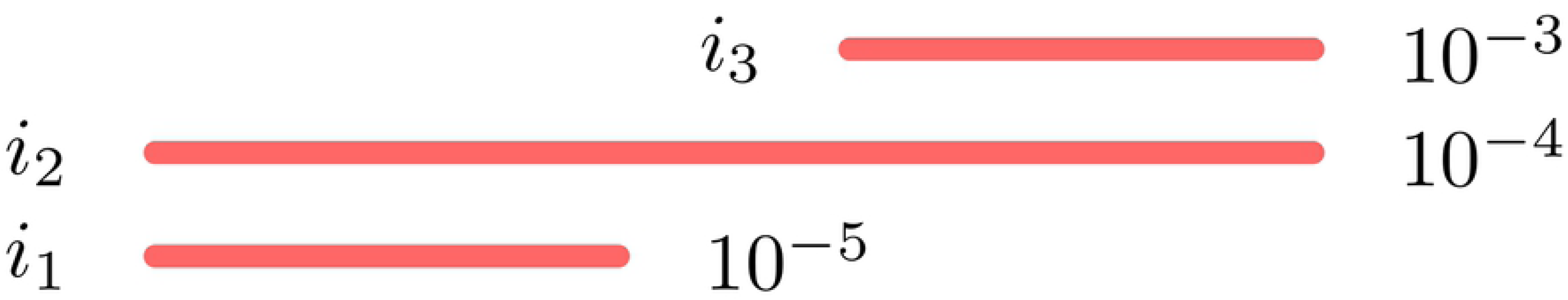
Merge step. Each interval *i*_1_ to *i*_3_, is associated to a p-value, written on the right of the interval. A naïve approach would discard *i*_2_ and *i*_3_ because they are dominated by *i*_1_ and *i*_2_ respectively. However, *i*_3_ may be an interesting interval, although the signal is not as strong as the signal of *i*_1_. We can notice that *i*_2_ both dominates (*i*_3_) and is dominated (by *i*_1_). Only this interval is discarded.

Our method only discards all the intervals that are both dominated, and dominate other intervals. When these intervals have been discarded, only undominated intervals remain, and they are given to the user (together with their p-values).

### Strategies

#### Preprocessing

Prior to the analysis of the data, the samples are first normalized using the CPM procedure, as done in edgeR [14].

Moreover, most of the strategy use a run-length encoding representation of the data, which is a compact way to store the expression of each nucleotide of the genome. This process is described in Fig 3.

**Fig 3.**
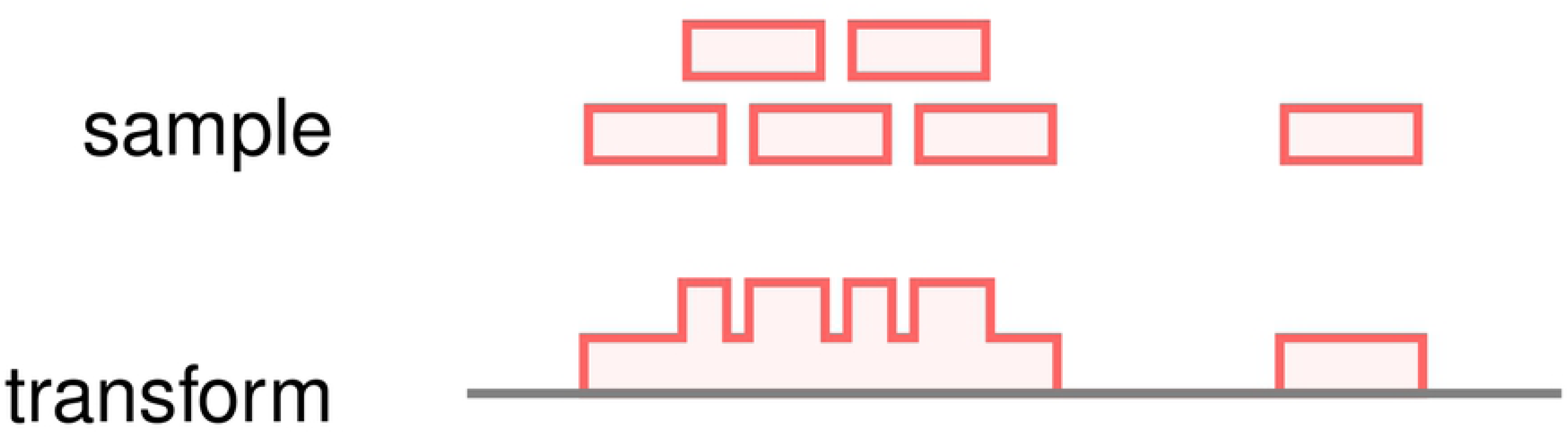
Transformation from mapped reads to run-length encoding. The reads themselves are lost, only the coverage is kept. For the sake of memory compactness, the coverage is stored as a vector of pairs (coverage, length) per chromosome.

#### Annotation

This step simply provides the intervals corresponding to the annotation file that is optionally given by the user. It can be a set of known miRNAs, siRNAs, piRNAs, or a combination thereof.

#### Naïve

The outline of the method is shown in Fig 4. This strategy compute the average of the expression in each condition. Then, the (log2) fold change of the expression is then computed. All the regions with a fold change greater than *n*, a parameter provided by the user, are kept as putative regions. The putative regions that are distant by no more than *d* nucleotides are then merged. However, we do not merge two regions if their log2 fold change have different signs. The remaining intervals are provided as candidate regions.

**Fig 4.**
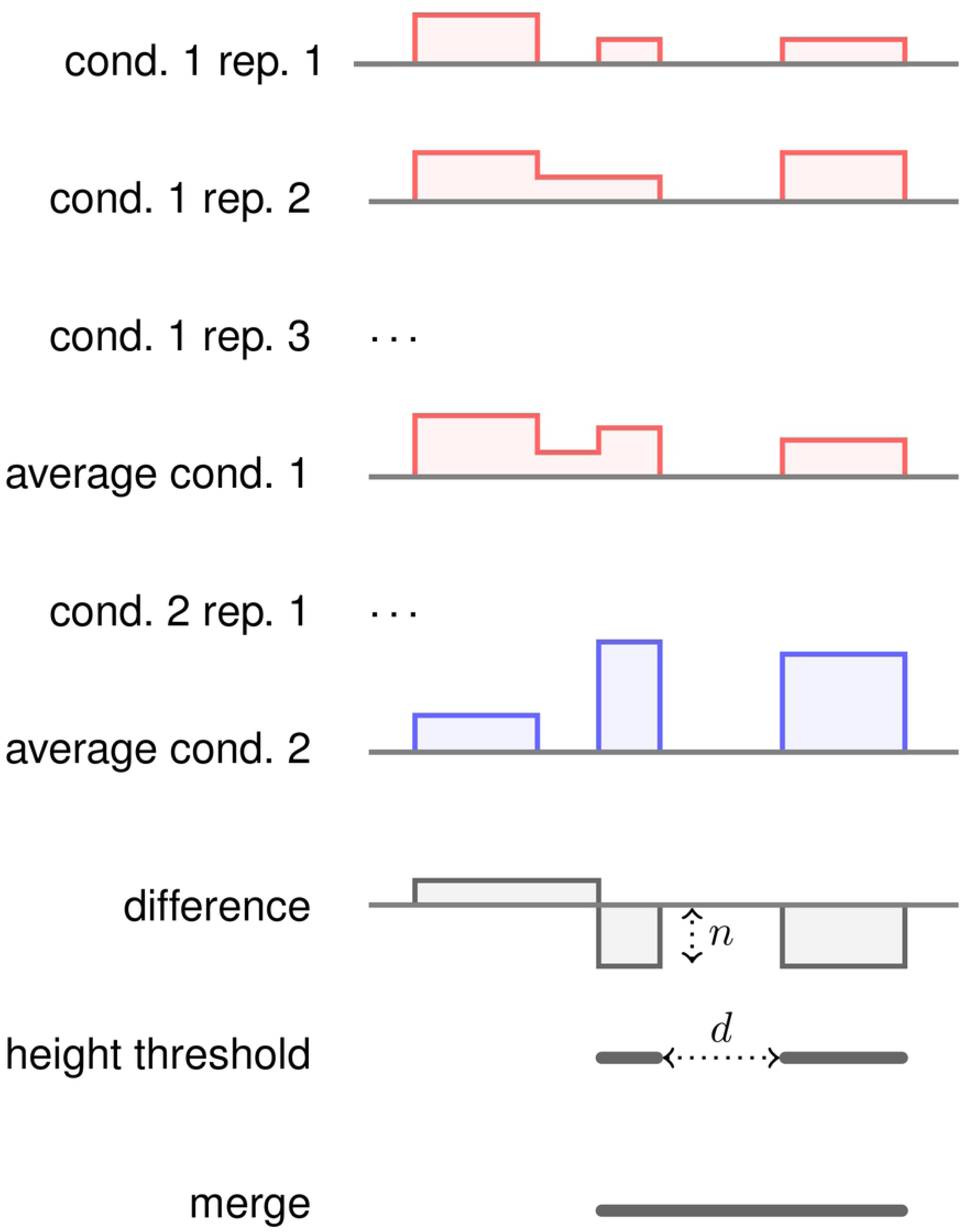
The naïve method. In a first step, the samples are averaged for each condition, then the (log2) fold change is computed. All those regions with a fold change not less than *n* are kept. Regions distant than no more than *d* base pairs are then merged, and given as output of the method.

#### HMM

We first form a matrix, where each line is a nucleotide, each column a sample, and each cell is the corresponding expression. This matrix is given to DESeq2. We then proceed to the standard DESeq2 workflow, and we compute an adjusted p-value for each nucleotide (see Fig 5).

**Fig 5.**
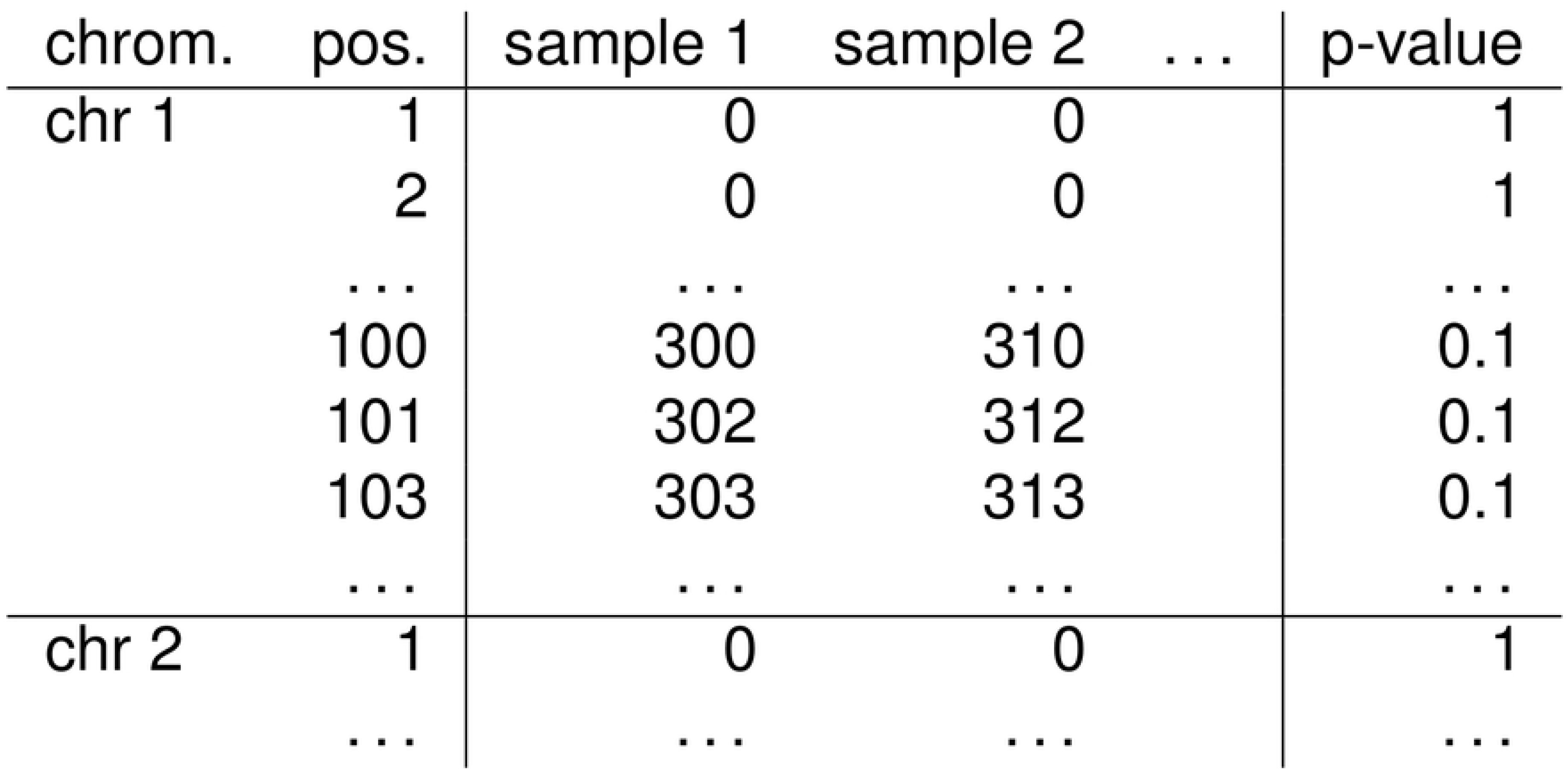
First step of the HMM method. The coverage of each nucleotide is computer for every condition. A p-value is produced for each position.

We then build an hidden Markov model (HMM) on each chromosome, where the first state is “differentially expressed”, and the second state is “not differentially expressed”, the observations are the p-values (see Fig 6). This HMM has been given sensible emission, transition, and starting probabilities values, but these parameters can be tuned by the user (S1 Appendix shows that the method does not seem sensitive to parameters). We then run the Viterbi algorithm, in order to have the most likely sequence of states. The regions that are most likely to be differentially expressed are given as output of the method.

**Fig 6.**
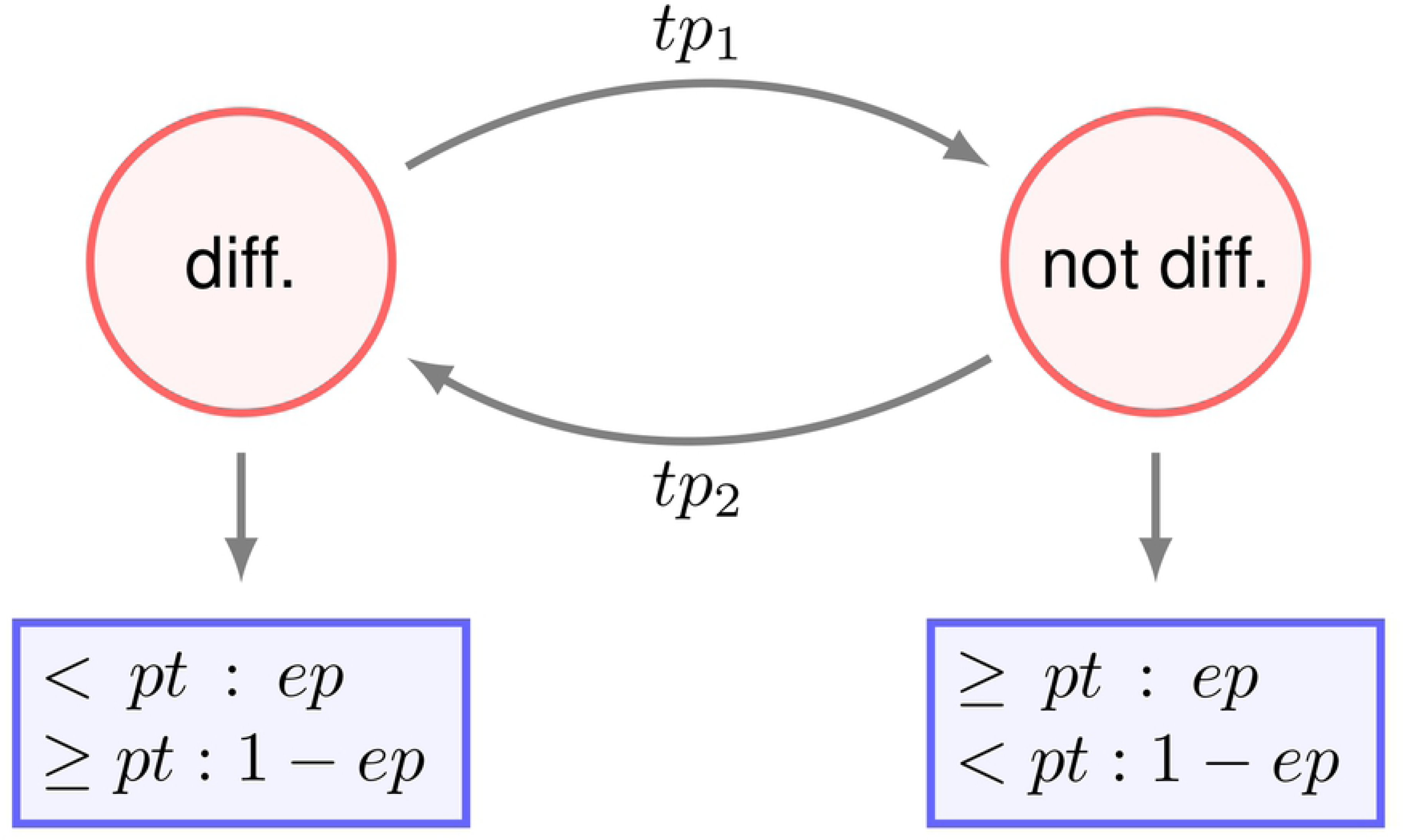
Second step of the HMM method. An HMM is run on each chromosome. The states are the red circles, and the emission probabilities are the blue rectangles. diff: the “differentially expressed” state. not diff: the “not differentially expressed” state. *tp*_1_ and *tp*_2_ are the transition probabilities. The emission of each state follows a binomial distribution. For instance, the diff. state emits a p-value less than *pt* with probability *ep*. All the parameters (*tp*_*i*_, *pt* and *ep*) are editable by the user.

In practice, the p-value is not computed for every nucleotide. Regions where the sum of the coverage is less than a threshold (editable by the user) are given a p-value of 1, because these poorly expressed regions are unlikely to provide a differentially expressed sRNA. In the HMM, all the regions with a p-value of 1 (the majority of the genome, because small RNA transcription is restricted to a minority of loci) are not stored and are assumed to have the default value. This significantly reduces the memory consumption. During the Viterbi algorithm, the most frequent state is “not differentially expressed”, and the most frequent p-value is 1. In this configuration, if the probability of “not differentially expressed” is significantly larger than the probability of the other state, we directly skip to the next nucleotide with a p-value < 1. Indeed, the difference of the probabilities of the “not differentially expressed” and “differentially expressed” states are, in this case, constant, and do not change the results the Viterbi algorithm.

#### Slice

The average (normalized) coverage of each condition is then computed. We then compute the (log2) fold change, and find irreducible regions (IRs), as presented in [15]. The method is presented in figure 7. Briefly, the method extracts all the regions where the fold change is above a threshold (given by the user). The IR method is simple and efficient way to merge such regions when they are not very far away, and the drop in fold changed is not too deep.

**Fig 7.**
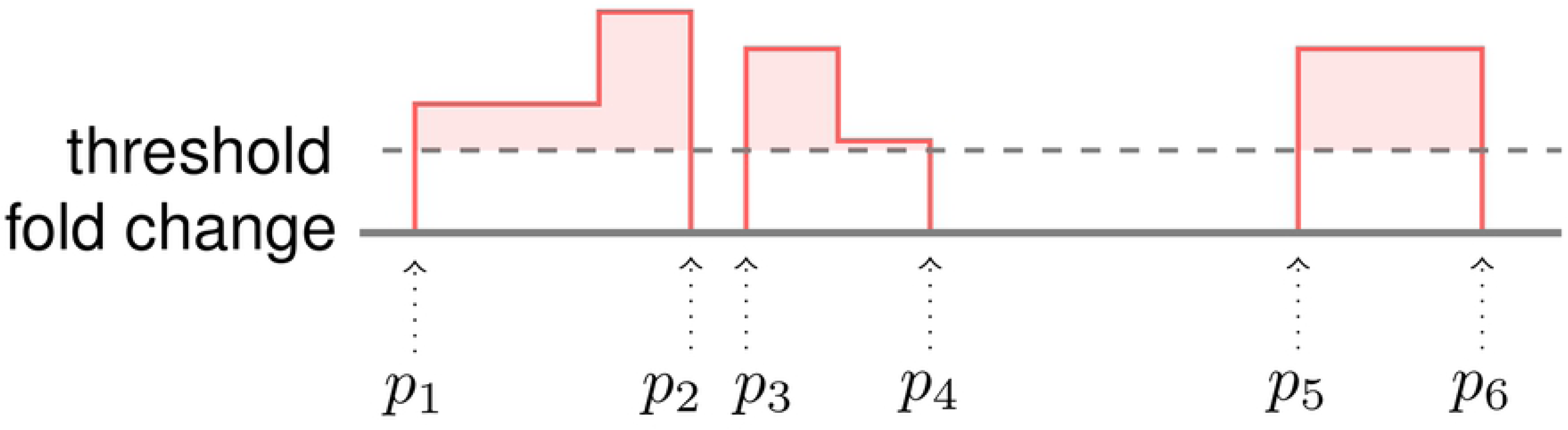
IR step. The (log2) fold change is printed in red, 0 is given in solid grey, and the dashed line is the (user given) threshold. Every region above the threshold is a putative differentially expressed region. A simple method could give three regions (between *p*_1_ and *p*_2_, between *p*_3_ and *p*_4_, and between *p*_5_ and *p*_6_). The IR method aims at merging close-by regions, with no additional parameter (contrary to the naïve method). Briefly, the method considers every interval (*p*_1_–*p*_6_, for instance), computes the area above the threshold (in light red), and divides it by the size (here, *p*_6_ –*p*_1_). We will call the mean area above the threshold MAAT. The method then considers all the positions, for instance *p*_3_, between *p*_1_ and *p*_6_. If the MAAT between *p*_1_ and *p*_3_, or the MAAT between *p*_3_ and *p*_6_, is not above the threshold, the interval is split. In the example, the MAAT between *p*_3_ and *p*_6_ is visibly less than the threshold, so the region is split at *p*_3_. However, the region between *p*_1_ and *p*_4_ is not split, since the MAATs between *p*_1_ and *p*_2_, *p*_1_ and *p*_3_, *p*_2_ and *p*_3_, *p*_2_ and *p*_3_ are all greater than the threshold.

In practice, the IR method can be very efficiently implemented. It simply requires a linear time algorithm, that considers all the points where the fold change intersects the threshold.

We also take care not to merge regions with positive log fold change, and regions with negative log fold change.

### Implementation of srnadiff

#### Example of use

srnadiff can be installed through R Bioconductor [16]. As every Bioconductor package, it has a dedicated Web page (https://bioconductor.org/packages/release/bioc/html/srnadiff.html) and contains an extensive description of the tool.

We provide here an introduction on how to use the package.

The information about the data set can be conveniently stored in a `data. frame`. This table should contain three columns, and each row describes a sample. The columns should refer to the name of the BAM files, the name of the sample, and the condition (e.g. `wild` type *vs* `mutant`).

If the table is stored into a file named `data`.`csv`, the minimal code to run srnadiff is:

~~~
1  library (srnadiff)
2  data           <- read .csv(“data. csv”)
3  bam Files      <- file .path(dir, data\$ FileName)
4  exp            <- srnadiffExp (bam Files = bamFiles,
5                                        sampleInfo = data,
6                                        annotReg = “annotation. gtf”)
7  exp            <- srnadiff(exp)
8  diffRegions    <- regions(exp, 0.05)
9  plotRegions(exp, diffRegions [1])
~~~

In this code, `annotationFile.gtf` is a GTF file that contains the known annotation. It is an optional parameter.

The `srnadiffExp` function reads the input data, and transforms the BAM files into run-length encoding data. It returns an object of class `srnadiffExp`.

The `srnadiff` function performs the main tasks of the package: segmentation, reconciliation, and computation of the p-values. The parameters that control the algorithms can be changed using this function. The `segMethod` parameter takes the list of the segmentation methods that should be used (default is “HMM” and “IR”). The `nThreads` parameter controls the number of threads used. The other fine-tuning parameters (such as the minimum sequencing depth, the minimum and maximum feature ranges, etc.) are stored into the `srnadiffDefaultParameters` object. This object can be changed as desired, and provided to the `srnadiff` method.

The `regions` function provides the differentially expressed regions, in a `GenomicRanges` object [17]. A minimum (adjusted) p-value can be provided as parameter.

The `plotRegions` function is a utility tool, which plots the coverage of the different samples around a region of interest (usually a prediction of srnadiff). This function accept a great number of parameters to customize the plot (visual aspect, other annotation, etc.)

### Benchmarking

We benchmarked srnadiff on three real, already published datasets, and a synthetic one. The published datasets encompass a variety of model organisms (*Homo Sapiens, Arabidopsis thaliana* and *Drosophila melanogaster*), protocols, and sequencing machines. All the publications provided a list of differentially expressed miRNAs, and we compared the different methods with this list of miRNAs.

srnadiff was run with no annotation, and an adjusted p-value threshold of 5%. We also run derfinder [13] on the same datasets, with a q-value of 10%. We used a third method, which first clusters the reads with ShortStack [12] (comparing several clustering methods is out of the scope of this article), quantified the expression of the regions found by ShortStack with featureCounts, tested for differential expression with DESeq2, and kept the regions with an adjusted p-value of at most 5%. We refer to this method as the *ShortStack* method. The reason why we chose a q-value of 10% for derfinder, instead of 5%, is that the statistics produced by derfinder is significantly more conservative, and it produces much less predicted regions than other approaches. For a fair comparison, we decided to lower the stringency for this tool.

We then wanted to know which regions found by a given method were also detected by an other method.

We also compared srnadiff with another straightforward method: we downloaded available annotations of different sRNA-producing loci, and followed the previously presented method: expression quantification, and test for differential expression. These regions can be considered as true positives. For the human dataset, miRNAs were taken from miRBase [3], tRNAs (in order to find tRFs) from GtRNAdb [18], piRNAs from piRBase [19], snoRNAs from Ensembl [20], and genes (to find possible degradation products), from Ensembl too. The cress data was extracted from the TAIR annotation file [21], and from the FlyBase annotation file [22] for the fly. We then compared these differentially expressed regions with the predicted regions. Here, we stated that the two predicted regions *A* and *B* were similar when at least 80% of *A* overlaps with *B*, or at least 80% of *B* overlaps with *A*. The reason is that some annotation (such as genes, or tRNAs which are substentially largers that tRFs) as not expected to be differentially expressed. Moreover, ShortStack provides also significantly larger differentially expressed regions than srnadiff or derfinder.

For each tool, we plotted different results. First, we provided the size and the adjusted p-value distributions of the regions found. Then, we provided the number of differentially expressed features (e.g. differentially expressed miRNAs, tRFs, etc.) which overlap a given region found. The aim here was to test whether a method would “merge” several potential candidates into a unique, longer, differentially expressed region. Then, we focused on the regions found by srnadiff and another tool (derfinder or ShortStack). For each such region, we compared its size, and its (adjusted) p-value found by each method.

The code used for the benchmarking, and the versions of the tools used, are given in S1 Appendix.

### Datasets

#### Preprocessing

Published data sets were downloaded from SRA [23] using the SRA Toolkit. We cleaned the data with fastx clipper (http://hannonlab.cshl.edu/fastx_toolkit/index.html), mapped them with bowtie [24] (because it was ranked favorably in a recent benchmark [25]).

#### Human Dataset

The first data set compared healthy cells *vs* tumor cells of lungs of smokers. This data set has been sequenced on a Illumina GA-IIx, and contains 6 replicates per condition, with about 26 millions reads per sample. It has been published by [4], and re-analyzed by [5]. Both papers analyzed the sRNA-Seq data of lung tumors compared to adjacent normal tissues.

#### *A. thaliana* dataset

This dataset evaluated the difference of expression of small RNAs in two different concentrations of CO_2_ in *A. thaliana* [26]. Each condition contained two replicates, with about 11 millions reads each.

#### *D. melanogaster* dataset

In this dataset, [27] sequenced small RNAs of young and aged *D. melanogaster* flies circulating in the hemolymph. Each condition contained 8 and 4 replicates, with an average of 13 millions reads each.

### Synthetic dataset

We also generated a synthetic dataset, extracted from the human genome. We first selected 1000 miRNAs from miRBase [3], and 1000 piRNAs from piRNABank [6]. We randomly selected 100 upregulated miRNAs (2-fold change), 100 downregulated miRNAs (2-fold change), and the same for the piRNAs. We randomly assigned a baseline expression following a power law (*k* = 1.5) on the miRNAs and the piRNAs, which reflected a low expression for most of the RNAs, and a few very expressed RNAs, as observed on our data. We generated 6 replicates per condition using the polyester package [28]. Obviously some RNAs may be differentially expressed, but with a very low expression, they are almost impossible to detect. As a results, we quantified the expression of the features with featureCounts [29], tested for differential expression with DESeq2 [30], and restricted to all the features with an adjusted p-value of at most 5%. We considered these regions as our “truth” dataset.

## Results

### Human dataset

Results can be found in Table 1.

**Table 1.** Comparison of several approaches with srnadiff, derfinder, and a clustering method. The second column gives the number of regions found by each method named in the first column. The three last columns give the number of common regions that where also found by another method (srnadiff, derfinder, and ShortStack). The next two lines give the number of regions found by the two articles. The last four lines are the features that are detected as differentially expressed using the direct *annotate-then-identify* method.

| source | # regions | srnadiff | derfinder | ShortStack |
| --- | --- | --- | --- | --- |
| [4] | 48 | 34 | 17 | 37 |
| [5] | 23 | 19 | 13 | 10 |
| srnadiff | 1968 | — | 255 | 579 |
| derfinder | 464 | 256 | — | 196 |
| ShortStack | 617 | 463 | 169 | — |
| miRNAs | 240 | 181 | 48 | 169 |
| tRFs | 95 | 52 | 24 | 58 |
| snoRNAs | 42 | 29 | 1 | 29 |
| piRNAs | 10 | 6 | 4 | 8 |
| genes | 772 | 292 | 117 | 151 |

The first article found 48 differentially expressed miRNAs, and the second, only 23. Only 5 miRNAs were common in both analyses. We first wondered whether we could also detect these differentially expressed miRNAs using our method. First, the expression of six miRNAs found by [4] could not be correctly estimated because they belonged to duplicated regions in the new assembly (and not in the assembly used in the paper). Second, a miRNA found by [5] was missed because it was considered as an outlier by DESeq2.

We then compared these results with srnadiff (run with no annotation, and an adjusted p-value threshold of 5%). srnadiff finds 1968 differentially expressed regions in total. It missed a few miRNAs, because of an adjusted p-value threshold effect: when the test for differential expression is performed on a few miRNAs (here, 48 and 23 respectively), the adjustment is not expected to change the p-values. However, srnadiff has much more candidates (a few thousands), that should be tested. As a consequence, the adjustment is much stronger in this case, and many miRNAs have an adjusted p-value which is (slightly) greater than 5%. It is a usual trade-off between sensitivity and specificity.

Then, we compared the results with derfinder, which finds 464 differentially expressed regions. Most of the regions found by derfinder are also found by srnadiff. srnadiff missed some regions, because of the adjusted p-value threshold effect. Of note, no region found by derfinder with adjusted p-value less than 10^−5^ was missed. However, srnadiff provided significantly more regions.

We then compared with the ShortStack method, which finds 617 differentially expressed regions. Results show that srnadiff misses several regions. The main reason is that ShortStack accepts very large regions that may have a low p-value on the whole region even if the difference point-wise is not significant. The ShortStack method also finds a few more miRNAs found by the articles. However, srnadiff finds, in general, more than thrice as many regions.

Then, we compared all the results with a “truth set” (see Methods), where we retrieved several annotations, performed differential expression, and kept the regions with an adjusted p-value of 5%. We found, for instance, 240 differentially expressed miRNAs, 95 differentially expressed tRFS, etc. srnadiff usually is the method that recovers the greatest number of regions, although ShortStack sometimes provides more. It found 181 of the 240 differentially expressed miRNAs, 52 of the 97 tRFs, etc. Again, most of the missed features were due to the adjusted p-value threshold effect.

Last, srnadiff discovered 1581 differentially expressed regions outside of known small RNA genes, and 809 differentially expressed regions outside of known small RNA genes and any Ensembl annotation.

#### *A. thaliana* dataset

Results can be found in Table 2.

**Table 2.** Results on the *A. thaliana* dataset.

| method | # regions | srnadiff | derfinder | ShortStack |
| --- | --- | --- | --- | --- |
| [26] | 27 | 12 | 12 | 11 |
| srnadiff | 10975 | — | 3489 | 7569 |
| derfinder | 4974 | 4245 | — | 3274 |
| ShortStack | 7287 | 467 | 112 | — |
| genes | 1455 | 796 | 472 | 759 |
| miRNAs | 29 | 29 | 24 | 29 |
| tRFs | 62 | 38 | 35 | 39 |
| snoRNAs | 3 | 0 | 0 | 3 |

We applied the same methodology as previously. Here, more than half of the miRNAs found in the article was not detected by any other method, mostly because of the p-value threshold effect.

Here again, srnadiff usually gives better results than any other tool, with the exception of the differentially expressed snoRNAs.

#### *D. melanogaster* dataset

Results can be found in Table 3.

**Table 3.** Results on the *D. melanogaster* dataset.

| method | # regions | srnadiff | derfinder | ShortStack |
| --- | --- | --- | --- | --- |
| [27] | 19 | 4 | 1 | 4 |
| srnadiff | 2069 | — | 164 | 1152 |
| derfinder | 229 | 164 | — | 201 |
| ShortStack | 1203 | 911 | 98 | — |
| genes | 721 | 447 | 60 | 304 |
| miRNAs | 33 | 22 | 7 | 16 |
| snoRNAs | 9 | 7 | 0 | 8 |
| tRFs | 13 | 7 | 0 | 6 |

Similar conclusion can be drawn from this data set.

### Synthetic reads

The “truth” set contains 44 differentially expressed features. The results of each tool is given in Table 4. Precisions and recall are defined as 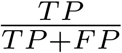 and 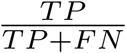 respectively, where *TP* is the number of true positives, *FP* is the number of false positives, and *FN* the number of false negatives. The *F*_1_ score is the harmonic mean of precision and recall, 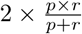, where *p* and *r* are precision and recall. srnadiff gives the best recall, ShortStack has the best precision, but srnadiff has a better combined *F*_1_ score. Oddly, derfinder did not give any predicted differentially expressed regions in this data set.

**Table 4.**
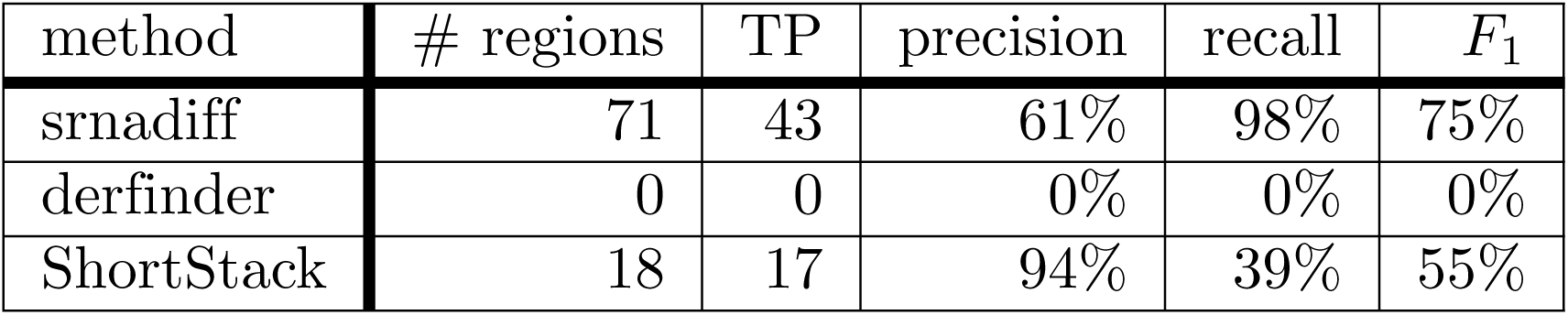
Results on the synthetic dataset. TP: true positives.

| method | # regions | TP | precision | recall | $F_1$ |
| --- | --- | --- | --- | --- | --- |
| srnadiff | 71 | 43 | 61% | 98% | 75% |
| derfinder | 0 | 0 | 0% | 0% | 0% |
| ShortStack | 18 | 17 | 94% | 39% | 55% |

### Time and memory usage

Table 5 provides time and memory usage of the tools used in the previous datasets.

**Table 5.** Time and memory usage of the tools. The first number in each cell is the time (in seconds), and the second is the memory (in Mb).

| method | human |  | *A. thaliana* |  | *D. melanogaster* |  | simulated |  |
| --- | --- | --- | --- | --- | --- | --- | --- | --- |
| srnadiff | 753 | 1824 | 134 | 2050 | 240 | 1518 | 240 | 1575 |
| derfinder | 491 | 2514 | 62 | 1327 | 59 | 1150 | 257 | 2066 |
| ShortStack | 1784 | 1304 | 232 | 901 | 582 | 2917 | 1274 | 689 |

Results clearly show that derfinder is the fastest tool, and ShortStack the slowest. However, srnadiff still provides results within 15 minutes. The reason of the difference between srnadiff and derfinder is that the former implements two methods, and thus processes the data twice. Second, derfinder uses bigWig files, whereas srnadiff readily uses BAM files (and internally converts them into a similar format, which is the bottleneck of the method). ShortStack is a Perl file, that requires significantly more time to process the data.

Concering the memory usage, derfinder is also the most efficient, and srnadiff usually the least efficient. However, all these computations fit in a standard computer.

All the computation has been performed on a personal computer running Linux Ubuntu 19.04, with Intel Xeon Processor E5-1650 v4 running 6 cores at 3.6 GHz and 32GB RAM.

Benchmarking the preprocessing steps (i.e. conversion from BAM to bigWig, and merging the BAM files for input of ShortStack) is not straightforward, because several pipelines are possible. We provide the usage we observed with deepTools [31] to convert BAM files to bigWig and samtools [32] to merge the BAM files in S1 Appendix.

Other benchmarking, available in S1 Appendix, show that changing srnadiff parameter does not significantly alter the results. We also show that time increases linearly with the input size. On the other hand, we also showed (see S1 Appendix) that the coverage does not have a dramatic influence on the results.

## Conclusion

In this paper, we propose a new method, called srnadiff, for the detection of differentially expressed small RNAs. The method offers several advantages. First, it can be applied to detect any type of small RNA: miRNAs, tRFs, siRNAs, etc. Second, it does not need any other knowledge on the studied small RNAs, such as an genome annotation, or a set of reference sequences. Moreover, results are comparable to *ad hoc* methods, which detect only a given type of small RNAs.

Our aim is to provide a simple tool that is able to extract all the information given by sRNA-Seq, not only restricting to miRNAs. We hope that srnadiff will make it possible to find new mechanisms involving understudied small RNAs.

Future directions for improvement include broaden the regions found, and providing new strategies to find differentially regions (besides the HMM and the IR methods).

## Supporting information

### S1 Appendix Additional Data

Supplementary figures, other benchmarking, code used, and tool versions are given in the Additional Data file.

## Acknowledgements

The authors thank Hervé Vaucheret, Stéphane Robin and Nathalie Vialaneix for fruitful discussions on the project.

